# Markov Chain Abstractions of Electrochemical Reaction-Diffusion in Synaptic Transmission for Neuromorphic Computing

**DOI:** 10.1101/2020.10.27.358390

**Authors:** Margot Wagner, Thomas M. Bartol, Terrence J. Sejnowski, Gert Cauwenberghs

## Abstract

Progress in computational neuroscience towards understanding brain function is challenged both by the complexity of molecular-scale electrochemical interactions at the level of individual neurons and synapses and the dimensionality of network dynamics across the brain covering a vast range of spatial and temporal scales. Our work abstracts an existing highly detailed, biophysically realistic 3D reaction-diffusion model of a chemical synapse to a compact internal state space representation that maps onto parallel neuromorphic hardware for efficient emulation at a very large scale and offers near-equivalence in input-output dynamics while preserving biologically interpretable tunable parameters.

## 1 INTRODUCTION

It has been known since the pioneering of computer architecture by John von Neumann that brains are far more effective and efficient in processing sensory information than digital computers, owing to the massively parallel distributed organization of neural circuits in the brain that tightly couple synaptic memory and computing at a fine grain scale (von Neumann, 1958). Modern day computers still follow the “von Neumann” architecture where computing and memory are kept separate, incurring severe penalties in computing bandwidth due to the bottleneck in data flow between centralized processing and vast memory. Moore’s law’s relentless scaling of semiconductor technology, with a doubling of integration density every two years, has allowed the von Neumann architecture to remain fundamentally unchanged since its advent. As the shrinking dimensions of transistors supporting the progression of Moore’s law are approaching fundamental limits, it has become essential to consider alternative novel computing architectures to meet increasing computational needs in this age of the deep learning revolution, which itself is driven by advances rooted in a deeper understanding of brain function (Sejnowski, 2020). At the forefront of this movement are neuromorphic systems, introduced by Carver Mead (1990) as a solution to these limitations. Neuromorphic engineering looks towards human brains as inspiration for hardware systems due to their highly efficient computational nature. The human brain is regarded as the pinnacle of efficient computing, operating at an estimated rate of 10^16^ complex operations per second while consuming less than 20 W of power (Churchland and Sejnowski, 1992; Cauwenberghs, 2013). Therefore, neuromorphic engineering looks to mimic the function and organization of neural structures using hybrid analog and digital systems. This is possible because there is significant overlap in the physics of computation between the brain and neuromorphic engineering (Figure 1). In both systems, information is carried in the form of charge, and, in hardware, neuronal membrane dynamics are represented using metal-oxide-semiconductor field-effect transistors (MOSFETs) (Mead, 1989). In the MOSFET sub-threshold region of operation, electrons and holes are the carriers of current between *n*- or *p*-type channels and behave akin to ions flowing through ion channels that mediate current across the neuronal cell membrane. Fundamentally, these hardware systems share analogous properties to their biological counterparts, including charge stochasticity, diffusion as the primary mechanism of carrier transport, and energy barriers modulated by gating voltage. Paired with Boltzmann distributions of charge, these systems are able to emulate current as an exponential function of the applied voltage, capturing the same biophysics underlying the neuronal dynamics (Mead, 1989; Broccard et al., 2017).

**Figure 1.**
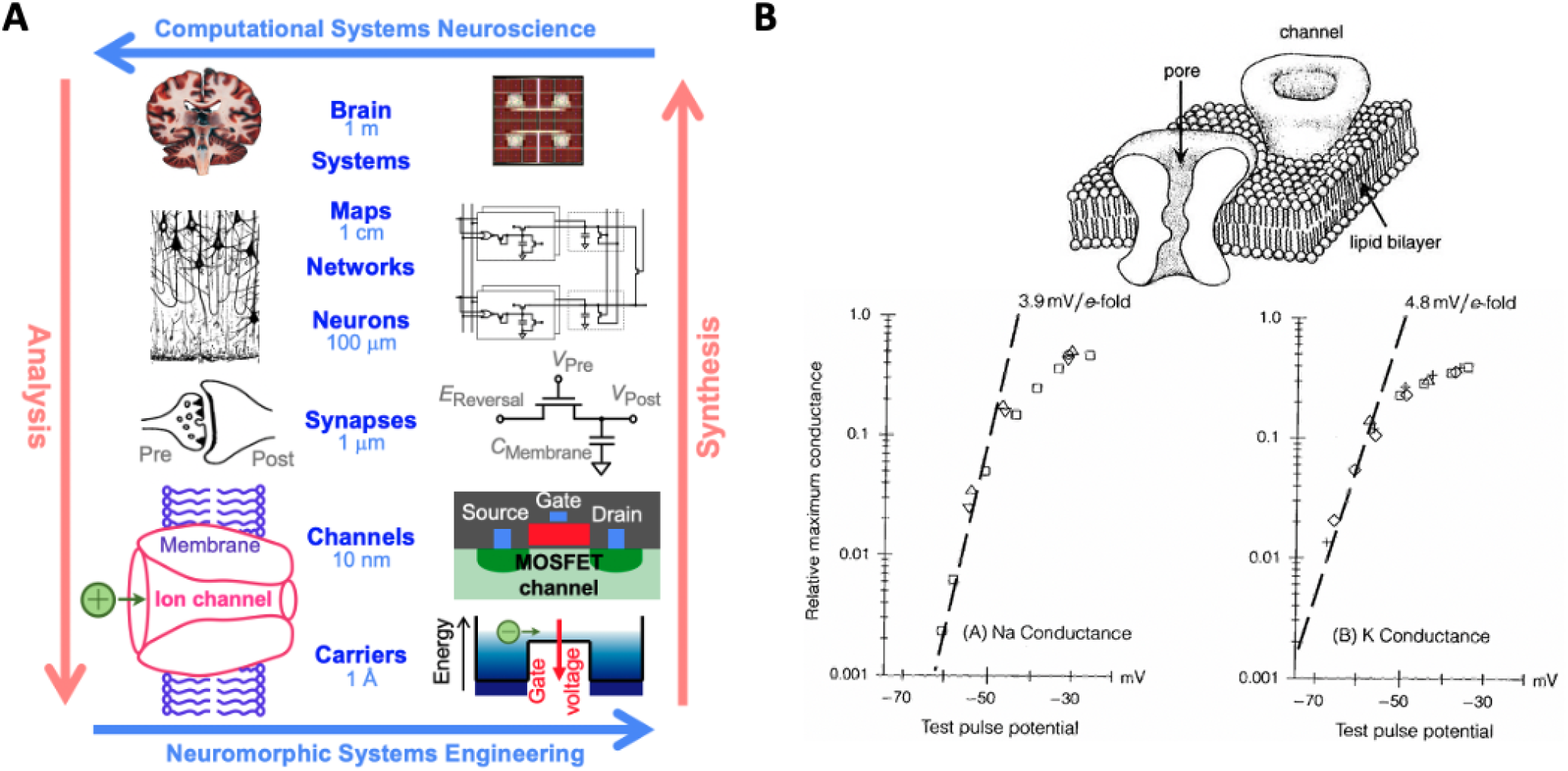
**(A)** Multiscale correspondence between the fields of computational neuroscience and neuromorphic engineering. Reproduced with permission from (Churchland and Sejnowski, 1992; Cauwenberghs, 2013) and **(B)** equivalence in Boltzmann thermodynamics between metal-oxide-semiconductor field-effect transistors (MOSFETs) and ion channels resulting in current as an exponential function of applied voltage in both systems for sodium *(bottom left)* and potassium *(bottom right)* (Hodgkin and Huxley, 1952; Mead, 1989).

Since the introduction of neuromorphic engineering, computational models of different complexity have been introduced to describe neuronal dynamics, typically ranging from more detailed and realistic conductance-based Hodgkin-Huxley models to simpler integrate-and-fire models allowing for better scalability. Synaptic connectivity between neurons is of primary concern in the field currently because synaptic strength and plasticity are fundamental to learning and memory in both biological and artificial representations of neural networks (Broccard et al., 2017; Indiveri et al., 2011). In neuromorphic architectures, synapses instantiate both computation and memory, and a new focus on compact electronic implementations of this computational memory has been emerging recently including the use of memristors (Boybat et al., 2018). Efficient representation of synapses is a crucial topic of concern as there are roughly 10^4^ synapses for each neuron, totalling approximately 10^16^ in the human brain. They are diverse in nature and have highly complex temporal and spatial dynamics, which further complicates their representations (Broccard et al., 2017). Currently, there is a push for efficient synaptic models while maintaining the intricate dynamical behavior exhibited biophysiologically. Current models include time-multiplexing synapses, analog bistable synapses, and binary synapses to name a few, but the need for scalable and dynamically complex models of synaptic function and transmission is still existent and critical (Broccard et al., 2017; Bartolozzi and Indiveri, 2007).

Modelling synapses is a challenging task due to their intricacy and sheer quantity. As noted above, there are an estimated 10^16^ synapses in the human brain. They vary in function and type, including both chemical and electrical synapses and exhibit behavior spanning multiple different temporal and spatial scales, as well as being highly stochastic in nature (van Rossum et al., 2003; Wang et al., 2020). Additionally, synaptic plasticity causes changes in synaptic strength over time associated with learning and memory. Synaptic transmission involves a multitude of mechanisms and molecular components, making simulations including all components not readily scalable. In order to capture the sophisticated dynamics of synapses in a scalable manner, abstractions have to be made according to the research problem in question. The stochastic nature of synapses also makes large scale simulations more complicated as modelling stochastic processes is typically more computationally demanding. It has been shown in multiple instances that the noise present due to the stochastic variability in synapses is highly integral to synaptic transmission, so this becomes an important feature to maintain (Malagon et al., 2016). For example, Moezzi et al. (2014) proved that models including ion channel noise in calcium channels paired with the existence of a presynaptic mechanism causing random delays in synaptic vesicle availability best capture the interspike interval behavior of auditory nerve fiber models. Additionally, multiple experimental works have found the existence of presynaptic vesicles that are released into the synaptic cleft with some probability (Castillo and Katz, 1954; Korn and Faber, 1991). There are multiple similar conclusions found in modelling and experimental results as recently discussed by McDonnell et al. (2016).

Synapses form the connections between neurons and the strength of these connections changes over time, forming the basis of learning and memory in both biological and artificial neural networks. The computations involved in accurately modelling the biophysics of synapses are complex due to the highly nonlinear nature of their dynamics, yet most of the neural network models in use today abstract synaptic strength to a single or small number of scalar values, tuned to a specific task. The learning rule for updating synaptic strength is then typically applied using abstractions of synaptic plasticity such as spike-time dependent plasticity and its causal extensions for scalable real-time hardware implementation (Pedroni et al., 2019). Physical constraints and limitations in VLSI implementations restrict the functional form of synaptic representation. In turn, these abstractions restrict the potential computing power of neuromorphic systems and restrain achievable benchmarks in approaching the functional flexibility, resilience, and efficiency of neural computation in the biological brain. Our work addresses the need for a more biophysically realistic model of the synapse with biologically tunable parameters to represent synaptic dynamics while offering a path towards efficient real-time implementation in neuromorphic hardware.

Synaptic transmission is dictated by a series of events initiated by presynaptic stimulation in the form of action potentials. An action potential causes membrane depolarization which leads to stochastic opening and closing of voltage-dependent calcium channels (VDCCs) lying on the presynaptic membrane and a resulting influx of calcium to the presynaptic terminal. Neurotransmitter release is modulated by calcium binding to calcium sensors near the neurotransmitter filled vesicles at the active zone, but calcium has other fates as it diffuses from the VDCCs. In addition to binding to the calcium sensors, it can bind to calbindin, which acts as a buffer, or it can be removed by plasma membrane calcium ATPase (PMCA) pumps. If enough calcium is able to bind to the calcium sensors, though, then neurotransmitters are released across the synaptic cleft and initiate downstream effects at the postsynaptic membrane (Bartol et al., 2015). This process of synaptic transmission is the basis of communication in the brain.

Abstracting this for computational efficiency, we created a series of Markov state transitions to realize the system with multiple internal states allowing for a biophysically tunable model of synaptic connectivity implementable in neuromorphic architectures. Markov models have a history of use as a stochastic discrete state alternative to Hodgkin-Huxley type formulations since their introduction (Hodgkin and Huxley, 1952; Armstrong, 1971; Colquhoun, 1973). Additional stochastic models have been introduced, including the Gillespie method (1977), which has been used to model neural channel noise (Gillespie, 1977; Skaugen and Walloe, 1979; Chow and White, 1996). Markov models have also found use in whole-cell models (Winslow et al., 1999). Further extensions utilize a particle model (Koch, 1999). The importance of the inclusion of stochasticity in ion channel behavior and synaptic transmission generally cannot be understated. Its inclusion has been demonstrated time and time again in experimental work and is thought to be integral in the form and function of synaptic transmission (McDonnell et al., 2016). This provides an additional complication in modelling synapses and has been handled at various different stages of transmission, including the stochastic models of vesicle release using probabilistically generated quantal components, stochastic models of transmitter diffusion, and stochastic models of receptors (Castillo and Katz, 1954; Bartol et al., 2015; van Rossum et al., 2003). These simulations are computationally expensive due to the high transition rates paired with the small number of transitions necessitating a small timepoint. Specifically, Markov models have shown to be an effective method of modelling ion channels but require high computational cost to effectively do so.

This paper looks to abstract the computationally complex and nonlinear nature of synaptic transmission dynamics in a manner that is efficient and readily scalable for implementation in neuromorphic silicon very large-scale integrated (VLSI) circuits. This is done by introducing an efficient stochastic sampling scheme within a Markov chain representation of the components integral to stochastic presynaptic quantal transmission.

## 2 MATERIALS AND METHODS

### 2.1 Markov Chain Models

The cascade of events from the action potential stimulus input to the presynaptic neurotransmitter release output can be equivalently modelled as a Markov chain to realize the system with multiple internal states instead of directly tracking all molecules and their kinetics in a computationally complex spatiotemporal 3D reaction-diffusion model. Each internal Markov state is assumed to be dependent solely on the state at the previous timepoint and is conditionally independent of all previous timepoints, simplifying simulations. Therefore, the fully biophysically complex system of synaptic transmission can be abstracted and sampled to create a Markov Chain Monte Carlo (MCMC) simulation which answers the same question of neurotransmitter release utilizing tunable biophysical parameters while providing scalability for implementation in neuromorphic architectures.

For any given stimulus input, the VDCCs are assigned transition probabilities between states based on a five-state kinetic model (Figure 2) found experimentally and validated computationally with four closed states and a single open state (Bischofberger et al., 2002; Church and Stanley, 1996; Bartol et al., 2015). Prior to the stimulus, all VDCCs begin in the initial closed state, C0, and the concentration of calcium in the presynaptic terminal is at steady-state. The transition probabilities are voltage dependent akin to a Hodgkin-Huxley model where 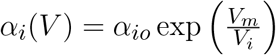 and similarly 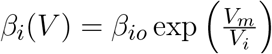 with parameter values from (Bischofberger et al., 2002). The number of open VDCCs at any given moment is used to determine the number of calcium entering the presynaptic terminal based on experimental I-V curves and the resulting I-V equation found in (Bischofberger et al., 2002) and used in (Bartol et al., 2015), which gives the value for *k*_*Ca*_. Calcium influx is captured by including transitions from the final closed VDCC state, C3, to the open VDCC state and an internal calcium generation. Using this, influx of calcium is modelled over the entire stimulus input due to the VDCCs opening.

**Figure 2.**
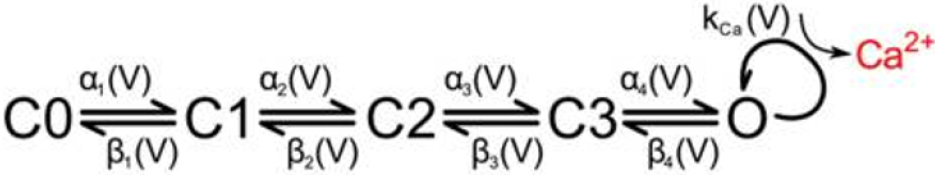
State diagram for voltage-dependent calcium channels and resulting calcium influx in the presynaptic membrane. Reproduced with permission from (Bartol et al., 2015).

Once calcium has entered the presynaptic terminal, much of it binds to calbindin, which acts as a buffer and primarily modulates the amount of calcium that is able to reach the calcium sensors at the active zone. The state transitions are reversible first-order reactions, thus transition probabilities are dependent on the free calcium in the system and updated as that amount changes over time. Calbindin has four binding sites, two of high affinity and two of medium affinity, leading to a nine-state calcium concentration-dependent kinetic model (Figure 3) (Nagerl et al., 2000). By modelling the binding and unbinding of calcium to calbindin as a loss or gain of free calcium respectively, calcium transients can also be elucidated.

**Figure 3.**
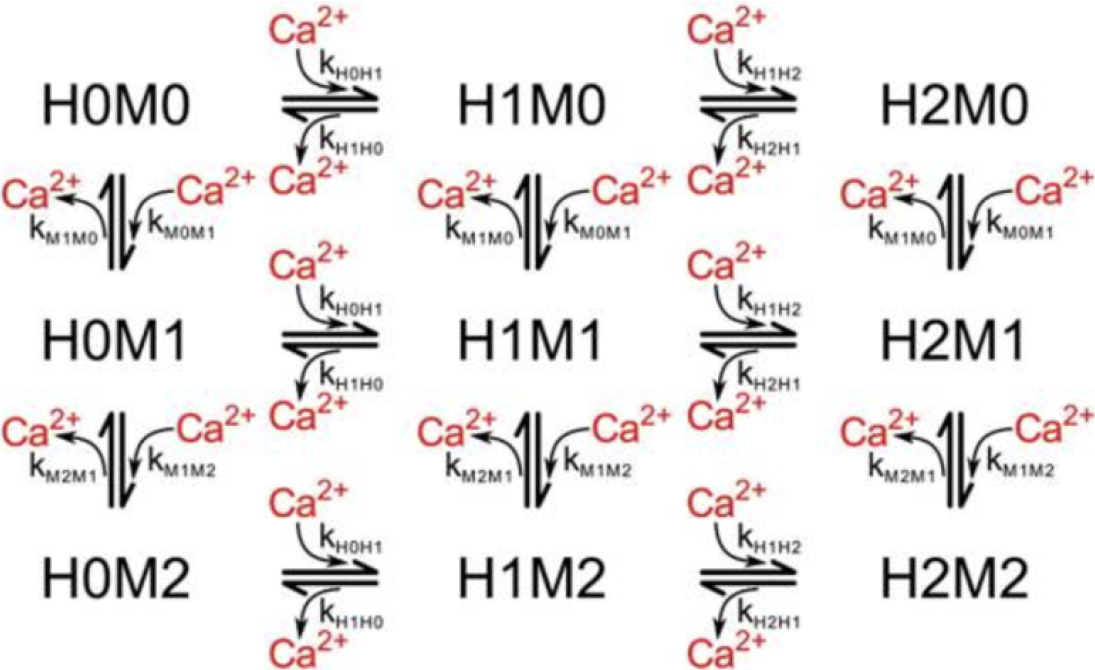
State diagram for calbindin binding where H*a*M*b* describes the *a*^*th*^ high-affinity binding state and the *b*^*th*^ medium-affinity binding state. Reproduced with permission from (Bartol et al., 2015).

Our Markov chain is a discrete-state chain in discrete time. Markov chains are modelled by a probability that the chain will move to another state given its current state and is conditionally independent of all previous timesteps. The probabilities are by nature only dependent on the current state of the Markov chain. The probability of the state of a molecule *X* can typically be predicted for a certain timepoint *t* + Δ*t* as some particular state *x*_*j*_ using the states at all previous timepoints from the start of the simulation, *t* = 0, to the timepoint just before that in question, *t*. For a Markov chain simulation solely dependent on the previous timepoint, it is possible to predict the probability that a molecule is in a given state, *x*_*j*_ at the timepoint *t* + Δ*t* using solely the state of the single timepoint just before, *X*_*t*_, which is known to be a particular state *x*_*i*_. Thus the probability of the molecule being in state *x*_*j*_ given that at the previous timepoint it was in state *x*_*i*_ is given as *P*_*ij*_. Succinctly, this is written as

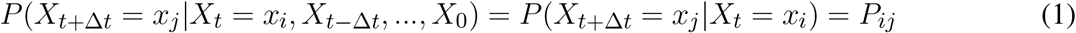

For state transitions, the probability of transitioning to an adjacent state is the transition rate inherent in the system (*k*_*ij*_ for the transition from state *i* to state *j*, and *k*_*ij*_ is not necessarily equal to *k*_*ji*_) times the timepoint, Δ*t*. In the case of calbindin transitions, this is further multiplied by the amount of free unbound calcium for forward reactions as it is a first-order reaction. For the VDCCs, the transition rates are the *α, β*, and *k*_*Ca*_. The probability that a molecule stays in its current state is the sum of the probabilities it transitions to an adjacent state subtracted from unity. For a multi-state system, this gives a transition probability matrix for the likelihood of transition from a given state at the current timepoint to any other state at the next timepoint. This matrix is sparse, with nonzero probabilities only for adjacent states to which a transition is possible. In the case of the five-state VDCC system, this gives the probability of a transition from state *i* to to state *j* as

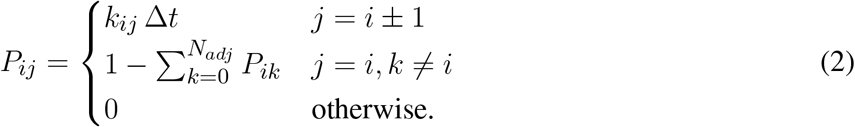

where transitions to adjacent states are given by the transition rate *k*_*ij*_ times the timepoint, Δ*t*; the probability of staying in the current state is the sum of probabilities of adjacent state transitions subtracted from unity, where *N*_*adj*_ is the number of possible adjacent states. The probability of transitioning to a non-adjacent state is set to zero.

Typically Markov state transitions are modelled via a discrete inverse transform method, where given a random variable *X*, the transition probabilities *P*_*ij*_ describe a partition of unity (Figure 4). Therefore, we can generate a random number uniformly, *R* ∼ ≤ *U* (0, 1) and map it onto discrete values of *X*. For example, in a two state system, *X*_*j*_ = 0 if *R ≤ P*_*i*0_ or *X*_*j*_ = 1 if *P*_*i*0_ *< R* ≤ *P*_*i*0_ + *P*_*i*1_ = 1. This involves searching the state space for the next state given the current state for each molecule in the system at each timepoint, which can be a slow process for systems with a large number of states and molecules.

**Figure 4.**
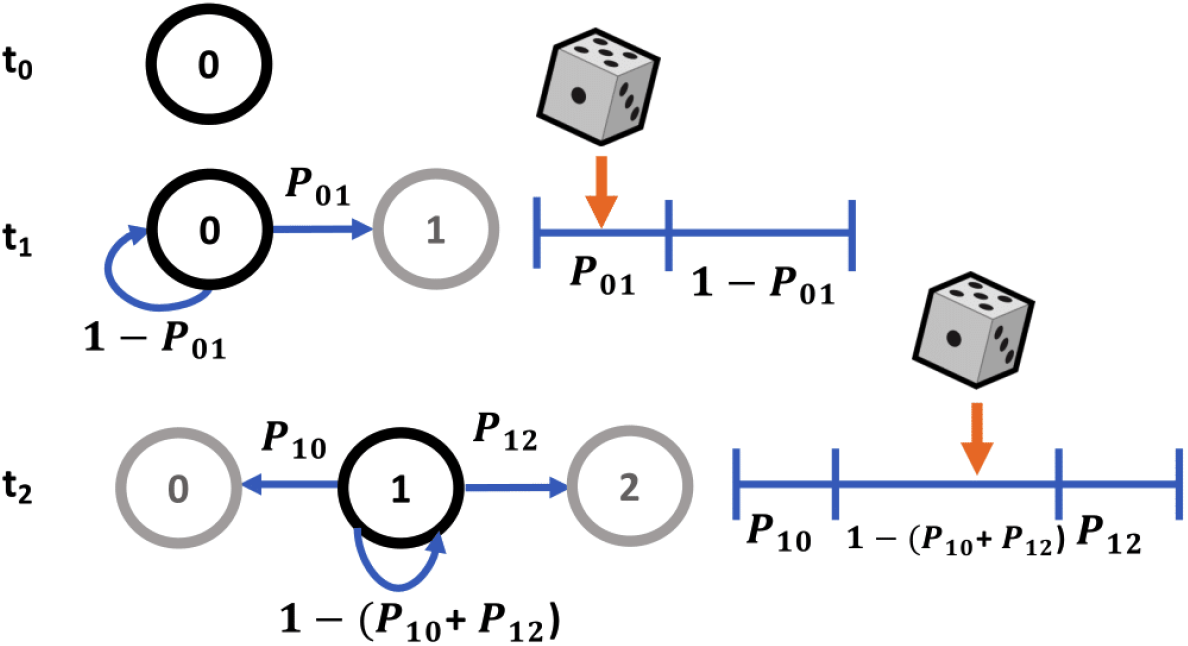
Markov sampling scheme for state transitions using partitions of unity.

Here we have implemented a more efficient MCMC sampling strategy involving sampling from a multinomial distribution. Therefore, instead of sampling from a uniform distribution for each of *n* molecules, we sample from a multinomial distribution once for each state, using *n* molecules as the number of experiments, where *X* ∼ Multi(*n, p*_1_, … *p*_*k*_). For simulations where the number of possible states is less than the number of molecules, this is a more efficient sampling strategy. Since we are particularly interested in the number of molecules in each state at each timepoint, this is an effective approach. Multinomial sampling thus describes the distribution of the *n* experiments across *k* possible outcomes each with a probability of *p*_*k*_, where *n*_*k*_ is the number of experiments falling into the *k*^*th*^ outcome following a probability mass function of

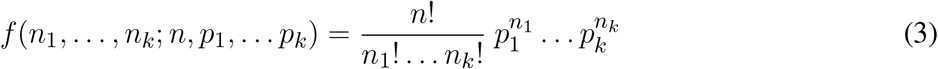

In our model, for each state *i*, we have an initial number of molecules in that state at a given timepoint *t*, or *n*_*i,t*_. As previously described, there exists a probability that the molecules will transition to any state at the next timepoint, including staying in the original state given by *P*_*ij*_. Thus, to determine the distribution of molecules *n*_*i,t*_ across all states at the next timepoint, we sample from a multinomial distribution according to

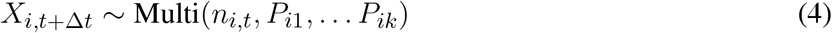

for *k* possible states. We do this sampling for each state at each timepoint and sum accordingly. This expedites computation by only requiring a single computation at each timepoint, sampling the distribution of all *n* molecules at once. Algorithm 1 highlights the pseudocode for this process.

#### Algorithm 1: Markov Multinomial Reaction Sampling

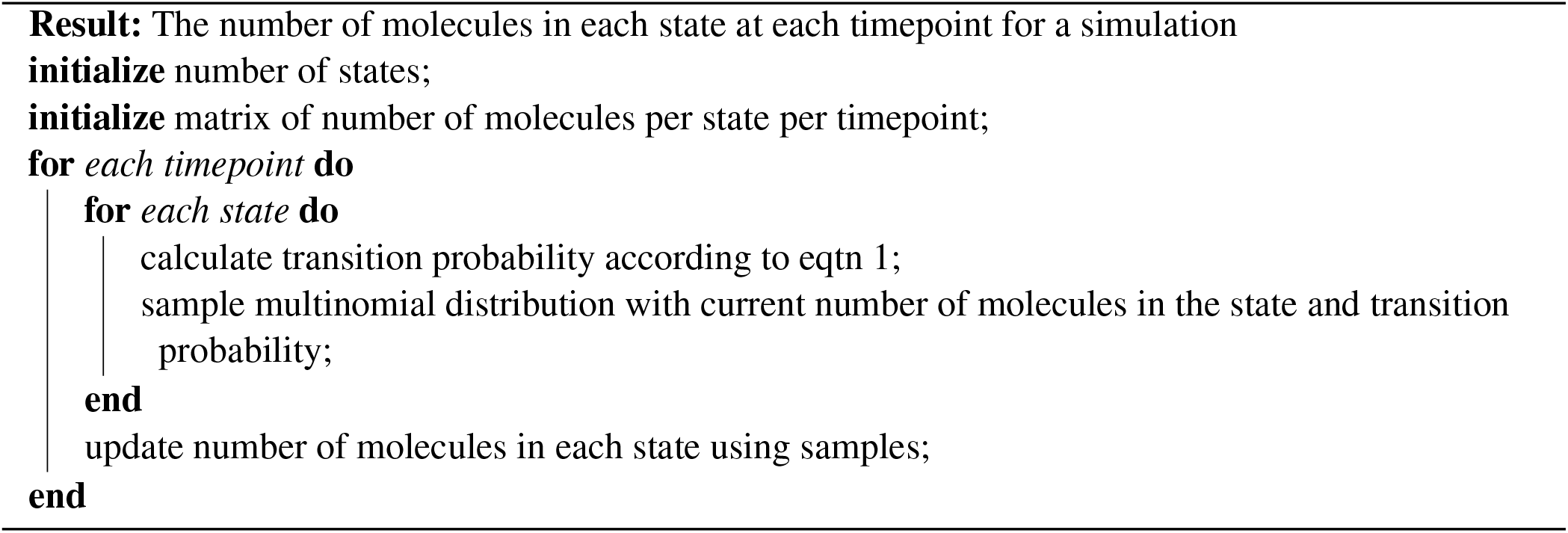

Markov simulations for the VDCCs were run for 65 VDCCs all starting in the closed state, C0. Calbindin molecules were initiated in the different binding states according to the steady-state concentration of calcium and at a baseline concentration of 4.5 10^*−*5^ M. All simulations were run for 10 ms with a timestep of 1 *μ*s. The simulations were repeated 1,000 times to obtain an average and standard deviation. Markov simulations were implemented using Python.

### 2.2 MCell Models

MCell is a modelling software that uses spatially realistic 3D geometries and Monte Carlo reaction-diffusion modelling algorithms, which allows for biophysically realistic simulations of high complexity as it specifically tracks the state of every molecules in space and time (Bartol et al., 2015). Due to the accuracy and specificity, it provides a ground truth for biological simulations but does so at the cost of computational complexity.

To validate and compare our Markov models of synaptic transmission, we built a biophysically realistic stochastic 3D reaction-diffusion system with all major components for presynaptic vesicle release variability in response to a stimulus input (Figure 5) based on the models of (Nadkarni et al., 2010) and (Bartol et al., 2015). The model includes realistic geometry for a CA3-CA1 *en passant* synapse focusing primarily on the presynaptic Schaffer collateral axon of a CA3 pyramidal cell found in the hippocampus with parameters set from experimental data (Nadkarni et al., 2010; Bartol et al., 2015). The CA3-CA1 synapse was chosen for the simulations as it is highly studied experimentally and is important for learning and memory. Furthermore, CA3-CA1 synapses are relatively small, containing one to two neurotransmitter release zones. Release from this region is also known to be highly stochastic in nature, necessitating the inclusion of stochasticity in biologically realistic models (Nadkarni et al., 2010). All kinetics and parameters match those used for the equivalent Markov models.

**Figure 5.**
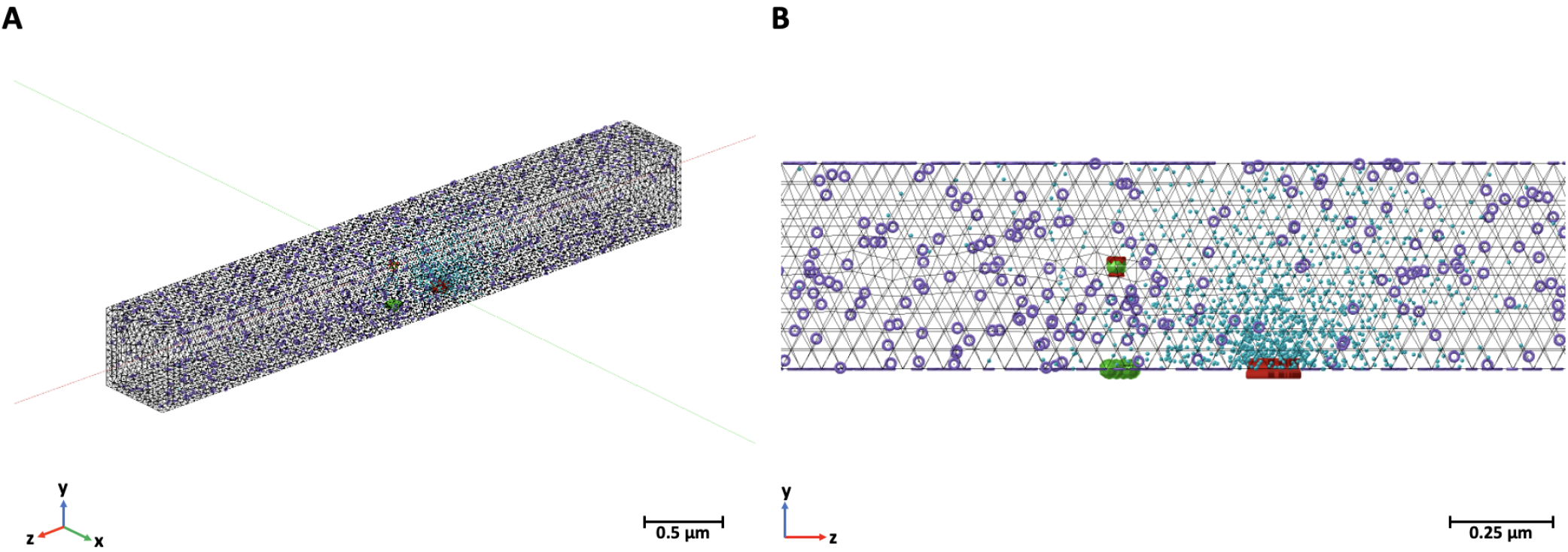
MCell model for synaptic transmission containing voltage-dependent calcium channels (red), calcium (blue), calcium sensors (green), and plasma membrane calcium ATPase pumps (purple). Calbindin not pictured here due to their large number. **(A)** Entire 0.5*μ*m by 0.5*μ*m by 4*μ*m box representing one vesicular release site in a Schaffer collateral axon in the CA3 region and **(B)** a close-up of the release site.

The MCell model includes the canonical presynaptic geometry for an average CA3-CA1 synaptic terminal as a rectangular box measuring 0.5*μ*m by 0.5*μ*m by 4*μ*m. This box captures the dynamics of a single synaptic active zone, referring to the region on the presynaptic membrane specialized for neurotransmitter release. Initially, the terminal contains the calbindin buffer, steady-state calcium concentration, PMCA pumps, VDCCs and calcium sensors modulating neurotransmitter release (Nadkarni et al., 2010). The detailed diffusion dynamics and kinetics of these systems are based on experimental data and have been discussed in further detail in (Bartol et al., 2015). The active zone is based on that of an average presynaptic active zone containing seven docked neurotransmitter vesicle release sites. The VDCCs, of type P/Q, are stationed at a biophysically realistic distance from the active zone. They transition states in response to the membrane depolarization. The location, number, and calcium conductance of the VDCCs is replicated from experimental data (Nadkarni et al., 2010). PMCAs are homogenously placed across the presynaptic membrane while calbindin molecules are in a uniform concentration within the volume. This is a flexible architecture that can respond to any stimulus input and allows for monitoring of the states of each molecule in the system. The MCell CA3-CA1 synaptic transmission models were originally created and validated in (Nadkarni et al., 2010) and (Bartol et al., 2015). To compare with the Markov models, we used the same single action potential stimulus.

MCell models were also run 1,000 times for 10 *ms* with a timestep of 1 *μs*.

## 3 RESULTS

### 3.1 Voltage-Dependent Calcium Channels

The efficient Markov chain implementation has strong agreement with the full MCell model in terms of the internal state transients in response to an external stimulus. The number of closed VDCCs (state C0) decreases over the duration of the stimulus (Figure 6A). The internal states (C1-C3) subsequently increase and decrease as the membrane voltage increases and the forward rates for the VDCCs increase (Figure 6B-D), leading to an exponential increase in the open VDCCs while the membrane depolarizes. Figure 6E shows the fraction of open VDCCs over time in response to the action potential, which controls the amount of calcium influx to the system. At the maximum membrane potential, almost all VDCCs are in the open state. As the membrane repolarizes, the reverse reaction rate constants increase, and the VDCCs close. This leads to another increase and decrease in the internal VDCC states as the receptors go from their open to resting closed state (C0).

**Figure 6.**
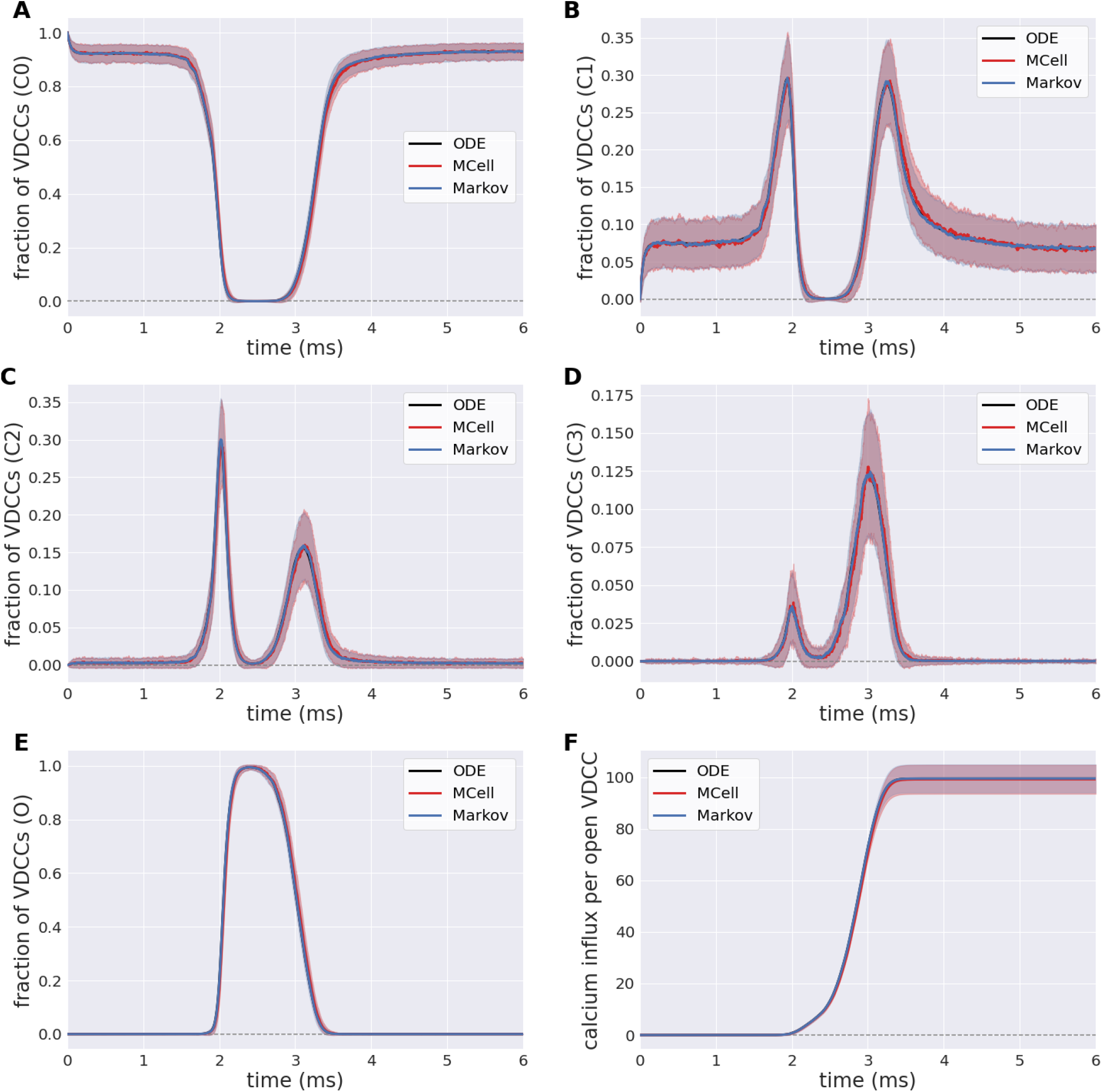
Fraction of voltage-dependent calcium channels (VDCCs) in each state: **(A)-(D)** Internal closed channel states, C0-C3, and **(E)** The open channel state, O. **(F)** Calcium influx through open VDCCs in response to action potential stimulus for stochastic MCell, Markov, and deterministic ODE forward Euler simulation.

In its open state, VDCCs allow for the probabilistic influx of calcium through the channels into the presynaptic bouton. This is exemplified in Figure 6F, where there is an increase in the calcium influx through the open VDCCs over the course of the stimulus. Again, there is strong agreement between the more computationally complex MCell model and the computationally efficient Markov equivalent model.

### 3.2 Calbindin Buffer

Simulations of homogeneous calcium and calbindin were run using the MCell, Markov and deterministic simulation schemes. In the presence of calcium, the forward binding reaction is heavily favored, and this is highlighted in Figure 7A where free calcium exponentially decreases. A similar transient is apparent for the unbound state of calbindin, as it quickly transitions to different stages of high and medium binding. Over the course of the simulation, all the free calcium is removed from the system, and calbindin states reach a new steady-state where there is still unbound calbindin. Similarly, the fully bound state, H2M2, rapidly increases and reaches a new steady state that is still only 1% of all calbindin. This is due to the high concentration of calbindin in the presynaptic bouton. Even once all the calcium is in a bound state, there is still plenty of unbound or partially bound calbindin remaining in the system. Calbindin acts as a strong buffer allowing for calcium storage and asynchronous neurotransmitter release, so this and slow unbinding rates become an important feature of calbindin. The rapid extent to which calcium binds to calbindin shows the impact of buffering on calcium’s ability to diffuse and bind to the calcium sensors in the active zone. The inclusion of calbindin at such high concentrations becomes a key feature of maintaining the relatively low release rates of neurotransmitters even in the presence of a stimulus.

**Figure 7.**
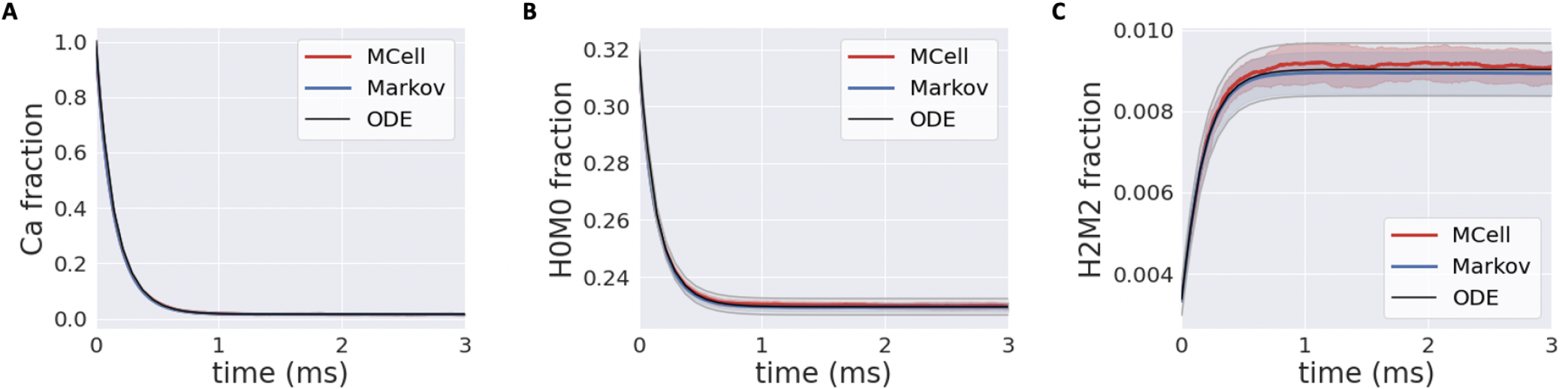
**(A)** Transients for homogeneous calcium-calbindin buffer binding in the presynaptic bouton for free calcium; **(B)** the unbound calbindin state, H0M0; and **(C)** the fully bound calbindin state, H2M2 in all simulation types.

### 3.3 Complexity Analysis

MCell uses a scheduler which allows for only making changes to the scheduled particles, though in the worst-case, this still scales with the total number of particles in the simulation, *n*, where *n*_*V DCC*_ is 65 and *n*_*calb*_ is 2.7 × 10^4^. It also scales with the length of the simulation, *t*, described by the number of time points for a discrete simulation. The simulations for the VDCC and calbindin both use 10k timepoints. At each timepoint, a particle can transition to any of its adjacent or branched states, *b*, which is similarly described by a fan-out factor in electronic implementation. From the VDCC kinetic model described in Figure 2, *b*_*V DCC*_ is 1-2 depending on the state while the calbindin kinetic model in Figure 3 gives *b*_*calb*_ of 2-4. The overall time complexity for MCell is 𝒪 (*bnt*). The classical Markov representation tracks every particle. It also searches through the space of each adjacent state for potential state transitions at each time point. Therefore, classical Markov implementation similarly results in an 𝒪 (*bnt*) time complexity, or 𝒪 (*nt* log_2_ *b*) at best for implementation with an efficient search algorithm. Both the multinomial Markov model and the Euler ODE implementation describe the system in terms of the number of molecules in each state leading to a dependence on the total number of states, *s ≥b*, rather than the total number of particles. The total number of states for VDCC is 5 (Figure 2) while the number of states for calbindin is 9 (Figure 3). Due to efficient sampling methods, the multinomial Markov method is independent of the number of adjacent states, leading to a time complexity of 𝒪 (*bst*) for both the multinomial Markov and Euler ODE methods. Thus, our stochastic multinomial Markov model is equally amenable to large scale simulations as the deterministic ODE method that is typically used in simulations involving more synapses.

The traditional Markov sampling model and the MCell representation store the molecular states in bits for each particle as well as the states adjacent to the current state, leading to a space complexity of 𝒪 (*bn* log_2_ *s*) The efficient Markov model and ODE solution both simply store the number of molecules represented by bits in each state at each timepoint as well as the branched states resulting in a space complexity of 𝒪 (*bs* log_2_ *n*). There exists a trade off here between the number of particles in each state compared to the number of states where one is stored directly and one is stored as an index. Thus, for simulations where the number of states is less than the number of particles, the multinomial Markov model is an efficient representation of the system, which is typically the case for biochemical simulation. MCell is more efficient with large state-space systems, but the number of states could be sparsified in a multinomial Markov representation by implementing dynamic instantiation and annihilation of states. Additionally, unseen or rarely seen states could be ignored by truncating based on probability of a particle being in that state. This would functionally decrease the number of states in the system allowing for use of the multinomial Markov simulation method.

### 3.4 Benchmarks

Runtime and total floating point operations were used as metrics for comparison between the simulation methods (Table 2). We also looked at the number of pseudorandom number generator calls (*n*_*P RNG*_) between the simulations as this provides a metric to elucidate the differences observed in execution time between the simulations. Here we compare MCell, the standard Markov model, and the multinomial Markov stochastic models. The deterministic Euler solution is included as well for a non-stochastic comparison. Again, it is valuable to note the importance of stochasticity in these models. Significant work has shown the necessity of stochasticity in models of synaptic transmission in order to match experimental work. It has been demonstrated that deterministic models at this scale generally underestimate quantal release as concentration fluctuations are not captured (van Rossum et al., 2003; McDonnell et al., 2016; Bartol et al., 2015). Thus while deterministic ODE models provide efficient simulation techniques, they are not able to capture the full complexity of the dynamics of synaptic transmission, hence motivating the need for an efficient stochastic model.

In the VDCC simulations, the multinomial sampling MCMC model has a runtime on the order of the forward Euler deterministic solution. The MCell and the standard Markov stochastic models exemplify a runtime an order of magnitude higher. The number of operations is also higher for the MCell and the standard Markov models compared to the multinomial Markov model. The standard Markov case generates a pseudorandom number for each molecule and each timestep, so *n*_*P RNG*_ is equivalent to the number of molecules multiplied by the number of timesteps, *n*_*V DCC*_*t*. In the multinomial Markov simulation, a pseudorandom number is generated for each occupied state and possible branching points at each timepoint, which gives (*bs*)_*V DCC*_*t* in the worst-case scenario. Therefore, *n*_*PRNG*_ is smaller for the multinomial case as long as (*bs*)_*V DCC*_ *< n*_*V DCC*_, which is always the case here.

For the calbindin model, the multinomial Markov method is again an order of magnitude faster than the MCell model although it is also an order of magnitude slower than the deterministic model. The standard Markov model is an order of magnitude slower than the multinomial model. The operations are also fewer for the multinomial case than the standard case. Again, the standard Markov case gives *n*_*P RNG*_ equal to *n*_*calb*_*t* while the multinomial Markov simulation is (*bs*)_*calb*_*t*. Again we see a smaller *n*_*P RNG*_ in the mulinomial case because (*bs*)_*calb*_ *< n*_*calb*_ even in the worst-case scenario where *b* is at its maximum value. Simulations are not currently optimized on hardware suggesting opportunities for further decreases in runtime. Overall, the multinomial Markov simulation provides a computationally efficient alternative to stochastic MCell simulations while maintaining the biological accuracy.

### 3.5 Neuromorphic Implementation

Thermodynamic foundations of neuromorphic engineering suggest direct biophysical implementation of populations of ion channels with individual stochastic opening and closing of gating variables driven by thermal noise fluctuations (Mead, 1989). So it seems only natural to consider implementations using stochastic ODEs describing the rates of reaction kinetics under additive white Gaussian noise (AWGN):

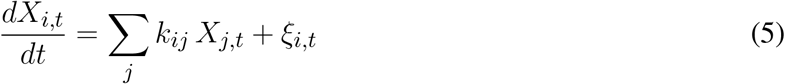

where *ξ*_*i,t*_ is normally distributed with zero mean and variance dependent on the magnitude of *X*_*i,t*_. Fully parallel, continuous-time analog implementation of reaction kinetic rate equations of the type (5) have been demonstrated in micropower integrated circuits, *e*.*g*., cytomorphic chips in BiCMOS integrated silicon technology (Woo et al., 2018). Abundant intrinsic noise present in these micropower cytomorphic circuits can serve as AWGN, although its magnitude is determined by thermal processes that are hard to control and other non-white Gaussian sources of intrinsic noise contribute strongly colored low-frequency spectra. Thus, discrete-time implementation of the ODEs (5) through Euler integration on a digital computer offers greater control over the shape and amplitude of the AWGN distribution, limited by the quality of pseudo-random number generation by deterministic algorithms.

Although purely digital algorithmic implementations go against foundational principles of neuromorphic engineering rooted in the physics of computation (Mead, 1989), the convenience of their programmability and reproducibility have made ODE-based digital emulation platforms such as Loihi a popular choice among more software-focused neuromorphic computer scientists (Davies et al., 2018). The computation involved in such discrete-time ODEs (5) can be performed at varying degrees of parallelism in custom or reconfigurable digital hardware, with the variables *X*_*i*_ being updated in sequence through time-multiplexing a single processing core in one extreme case, or all *X*_*i*_ updated in parallel with dedicated processing elements for each in the other extreme case. Ultimately in practice, the energy efficiency is relatively independent of the compute implementation, and depends more critically on the available memory bandwidth in accessing the rate parameters defining network connectivity (Pedroni et al., 2020). In essence, discrete-time Euler-integration ODE implementation of (5) amounts to sampling from a normal distribution

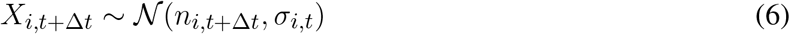

with mean and standard deviation

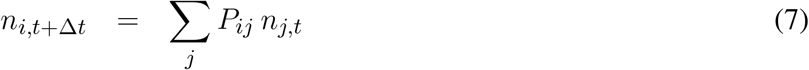

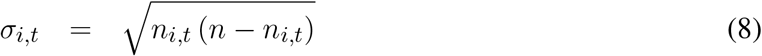

incurring computational complexity 𝒪 (*bst*) (Sec. 3.3).

More fundamentally, the main disadvantage of implementing stochastic ODEs (5) or their discrete-time digital versions (6) is that they are primarily based on the Central Limit Theorem for very large number of variables, *n* → ∞. As such, they have limited accuracy in approximating the reaction kinetics in systems with smaller numbers of molecular variables. While one may be tempted to assume that molecules are always excessively abundant, this is not typically the case since reactions are rate limited by the least abundant of reagents. Low numbers in molecular dynamics are prevalent in biologically relevant settings, giving rise to significant amounts of biological noise that are critical in neural dynamics, *e*.*g*.the highly stochastic quantal release of neurotransmitter in synaptic transmission. Thus, there is need for a mathematical description of stochastic synaptic transmission dynamics able to capture the accuracy in simulations with relatively small numbers of variables. Here we have shown that our multinomial Markov alternative, which directly samples the variables from the multinomial distribution (4) rather than the limiting normal distribution (6), produces accurate results for any value of *n* while offering nearly identical implementation complexity 𝒪 (*bst*) (Sec. 3.3). Hence we see the Markov chain abstractions of reaction kinetics not only as a means to approach biophysical realism in modeling molecular cellular dynamics without molecular-scale representation, but also as a means towards efficient neuromorphic hardware without biophysical compromise. The key point is that the computational complexity of implementing our multinomial Markov model is essentially identical to that of stochastic ODEs (see Table 1), whether in software executing serially on a von Neumann programmable digital computer or in massively parallel digital or analog hardware. Hence, the neuromorphic circuit designer tasked to implement brain-inspired models of information processing faces an easy choice: more bio-realistic models that account for detailed stochasticity in reaction kinetics incur the same resource utilization and energy costs, and use similar design principles, as their stochastic ODE approximations.

**Table 1.**
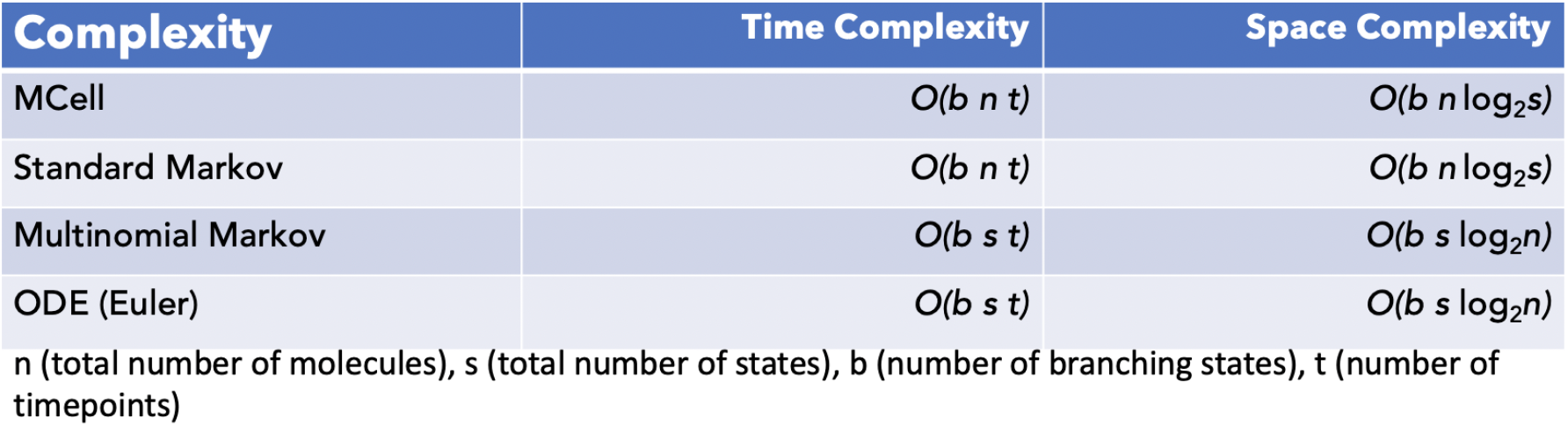
Space and time complexity for the various simulation strategies.

**Table 2.**
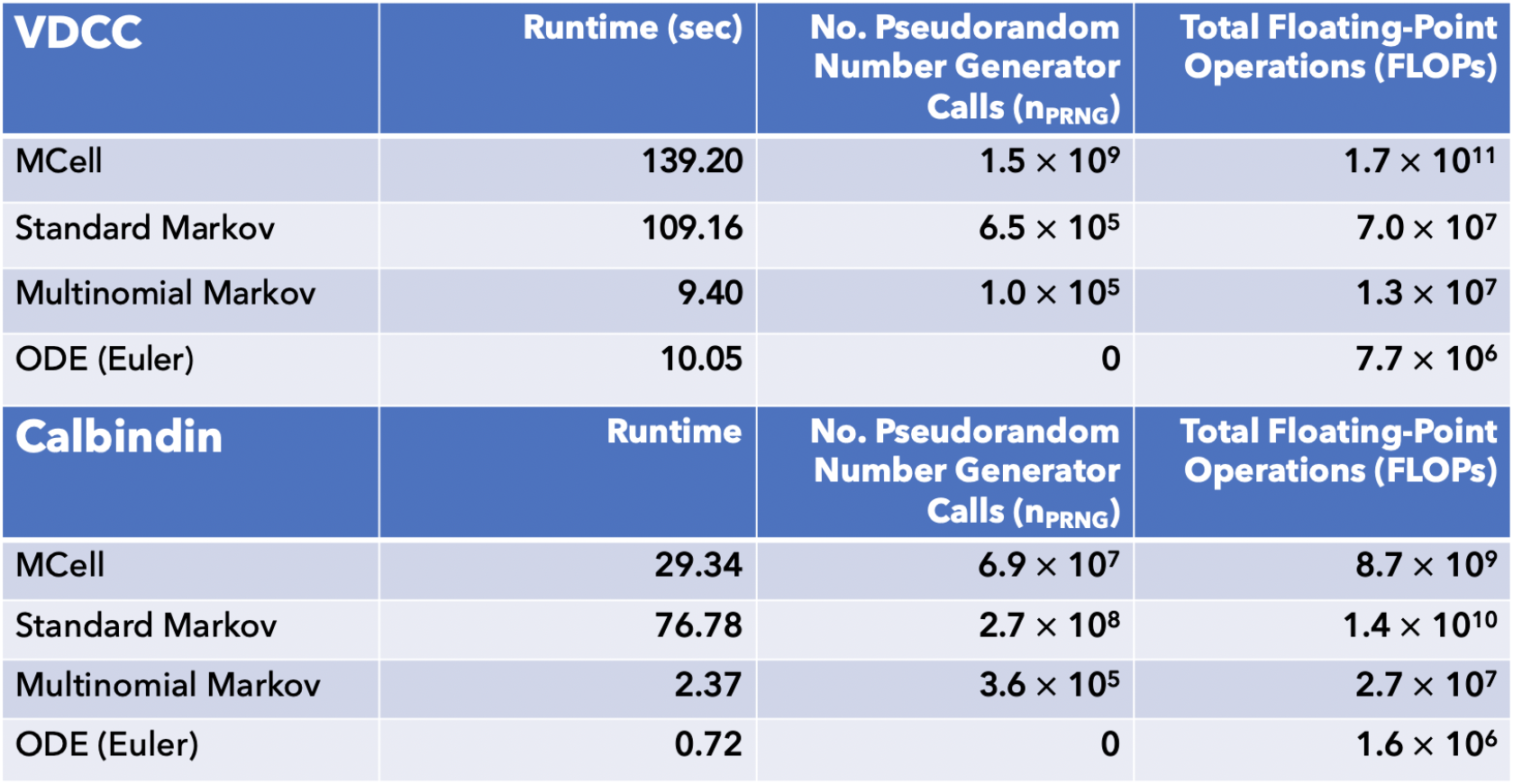
Benchmarks for different simulation types for both voltage-dependent calcium channel and calbindin binding simulations.

In addition to field-programmable gate array (FPGA) reconfigurable (Pedroni et al., 2020) or custom-integrated neuromorphic programmable (Davies et al., 2018) instantiations in digital hardware, we envision physically neuromorphic instantiations in micropower analog continuous-time compute-in-memory hardware that obviate sampling from posterior distributions and directly implement Markov state transitions through parallel implementation of sum-product rules with self-normalizing probabilities (Chakrabartty and Cauwenberghs, 2004, 2005), at throughput density and energy efficiency that are orders of magnitude higher than today’s most advanced general-programmable computational platforms.

## 4 DISCUSSION

The goal of this work was to create a more computationally efficient model of biologically realistic synaptic transmission for use in large-scale neuromorphic systems. We created a multinomial MCMC sampling strategy for capturing the internal states of vital molecules in the system in response to stimulus where transition probabilities could be voltage- or concentration-dependent, and the next timestep could be predicted solely using the current timestep. This scheme was implemented to capture the dynamics of the stochastic opening and closing of VDCCs through multiple internal states as well as the resulting calcium influx into the presynaptic bouton through the open VDCCs. Once calcium has entered the presynaptic terminal, we also simulated calcium binding to the calbindin buffer which modulates calcium levels in the bouton, directly impacting the amount of calcium that reaches the calcium sensors in the active zone. This amount impacts the neurotransmitter release from the presynaptic side and the resulting effects on the postsynaptic side.

All simulations were modelled using the multinomial Markov sampling method as well as a typical Markov sampling method and compared to highly detailed 3D geometric stochastic reaction-diffusion simulations done using MCell. The Markov simulations show agreement with the MCell simulations for the system dynamics including the number of open VDCCs and calcium influx in response to an action potential stimulus as well as the binding of calcium to the calbindin buffer. Differences are observed from the deterministic solution to the stochastic simulations implying the importance of stochasticity in these simulations to capture more biologically-realistic systems.

Exemplified by runtime and total number of operations, the multinomial MCMC method of simulations was shown to be more efficient than the standard Markov model while also being faster than the MCell equivalents. This is hopeful for scaling these biologically-realistic models to large-scale systems while maintaining biological tunability.

Next steps involve modelling the remaining kinetics in a similar fashion including the binding and removal of calcium by the plasma membrane calcium (PMCA) pumps as well as binding to the calcium sensors. In addition, to capture the diffusion of calcium through the presynaptic terminal but specifically to the calcium sensors at the active zone, a diffusive kernel must be included to the system. Upon inclusion of these elements, the entire process from stimulus to neurotransmitter release can be captured as a series of Markov chains leading to powerful implications for synaptic transmission modelling. The whole synapse can be included as well with the inclusion of a diffusive kernel across the synaptic cleft as well as downstream effects on the postsynaptic side, of which many mirror similar kinetics and dynamics as the presynaptic side leading to a natural extension of this modelling framework. The resulting system would be a biologically tunable model of synaptic transmission for any stimulus input in a highly efficient manner. This opens the door for large-scale implementations of synaptic transmission and learning readily implementable into neuromorphic architectures with strong biological realism.

Through the utilization of Markov-based abstractions applied to biophysically realistic 3D reaction-diffusion models of a chemical synapse, we have created a compact and efficient internal state space representation of synaptic transmission. This is in response to the challenge presented by the high dimensionality and complex nature of molecular-scale interactions in synapses and across scales making implementation in very large-scale systems previously unattainable. The model is directly amenable to efficient emulation in parallel neuromorphic hardware systems while maintaining biophysically relevant and interpretable parameters that are readily tunable. This opens the door towards neuromorphic circuits and systems on very large scale that strike a greater balance between integration density and biophysical accuracy in modelling neural function at the whole-brain level.

